# Enhanced Salt Tolerance in *Synechocystis* sp. PCC 6803 Through Adaptive Evolution: Mechanisms and Applications for Environmental Bioremediation

**DOI:** 10.1101/2024.08.29.610226

**Authors:** Xiaofei Zhu, Rongsong Zou, Dailin Liu, Jing Liu, Xuejing Wu, Lei Chen, Tao Sun, Weiwen Zhang

**Author notes:** These two authors contributed equally to this paper (Xiaofei Zhu, Rongsong Zou). To whom all correspondence should be addressed: Dr. Tao Sun, Center for Biosafety Research and Strategy, Tianjin University, Tianjin 30072, P.R. China., <u>Prof. Dr. Lei Chen</u>, Laboratory of Synthetic Microbiology, School of Chemical Engineering & Technology Tianjin University, Tianjin 300072, P.R. China.

## Abstract

Salt stress is common in natural environments, where elevated salt levels in brackish water and saline soil can hinder the growth of organisms, thereby exacerbating environmental challenges. Developing salt-tolerant organisms not only uncovers novel mechanisms of salt tolerance but also lays the groundwork for managing and utilizing saline environments. Cyanobacteria, which are widely distributed in hydrosphere and soil, serve as ideal models for studying salt stress. In this study, the model cyanobacterium *Synechocystis* sp. PCC 6803 was selected, whose salt (NaCl) tolerance improved from 4.0% to 6.5% (m/v) through adaptive laboratory evolution. Genome re-sequencing and mutant analysis identified six key genes associated with salt tolerance. Notably, the deletion of *slr1670*, which encodes glycerol glucoside hydrolase, improved the strain’s salt tolerance. In addition, *slr1753* encodes a membrane protein that may enhance salt tolerance by facilitating ion transport to the extracellular space. Further analysis revealed that overexpression of *slr1753* significantly accumulates Na^+^ on the cell surface, enabling effective seawater treatment using the engineered strain, resulting in a 6.35% reduction of Na^+^ in the seawater. Moreover, the adapted bacteria can be used for the remediation of saline soil samples, leading to a 184.2% and 43.8% increase in the germination rate and average height of *Brassica rapa chinensis*, respectively, along with a 25.3% rise in total organic carbon content and reductions in both total salt content by 1.82% and pH by 1.91% in soil. This study provides novel insights into salt tolerance mechanisms and the bioremediation of high-salinity environments.

**Graphical abstract:** 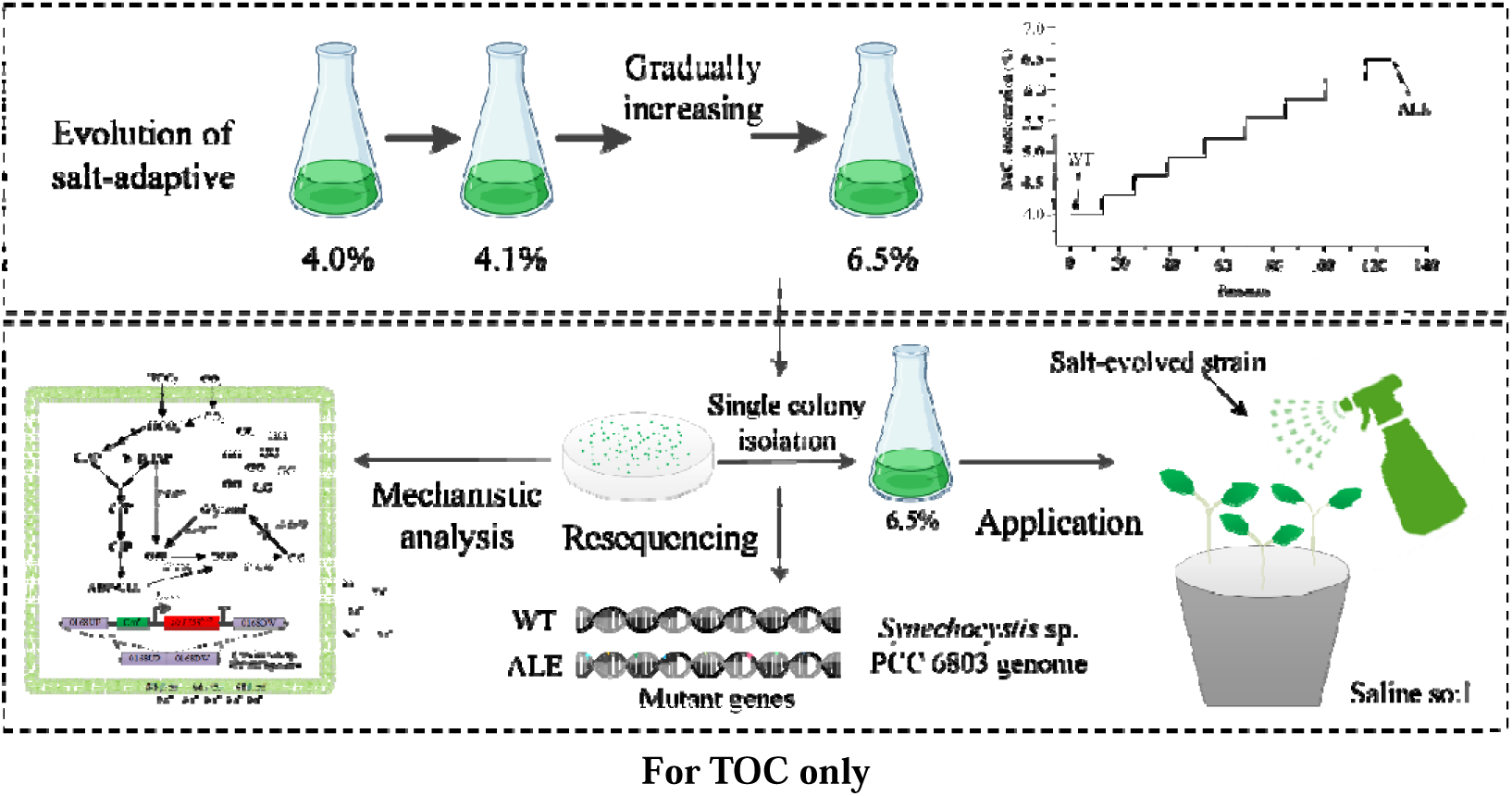

## Introduction

Salt stress caused by saline-alkali soil and brackish water, poses a significant environmental challenge with global implications for agricultural productivity and ecosystem health^[1]^. The presence of excess salt in soils or irrigation water has detrimental effects on plant growth, leading to substantial reductions in food production, particularly for major crops like rice and wheat^[2,3]^. Presently, the global extent of salt-affected land is estimated at approximately 1.1 × 10^9^ hectares^[4]^, encompassing 37.0 million hectares in China^[5]^, 5.7 million hectares in the United States^[6]^, and 67.2 million hectares in India^[7]^ among others. Furthermore, a substantial portion of arable land is undergoing salinization as a result of human activities such as deforestation, excessive irrigation, and other factors linked to climate change^[8]^. These include saline water encroachment into coastal regions due to rising sea levels and an increased incidence of storms^[9]^. Cyanobacteria, the earliest microorganisms on Earth capable of oxygen-producing photosynthesis^[10]^, have colonized all light-exposed habitats and play a pivotal role in global ecosystems^[11]^. Cyanobacteria are of significant interest due to their photosynthetic capabilities and stress resistance^[12,13]^. They contribute nutrients to the soil through nitrogen fixation, organic matter accumulation, and soil structure improvement^[14,15]^. It is well-documented that certain cyanobacteria establish symbiotic relationships with plants, fixing atmospheric nitrogen, promoting organic matter accumulation, and enhancing soil fertility^[16]^. Moreover, they secrete extracellular polysaccharides to bolster soil structure and water retention capacity^[17]^. Therefore, investigating the salt tolerance mechanisms of cyanobacteria is crucial for identifying salt-tolerant hosts not only for coping with saline soil environments but also for enhancing such environments.

To fight against salt stress, the physiological responses and molecular regulatory mechanisms of cyanobacteria have been extensively investigated^[18]^.Detailly, the response mechanisms include: i) Synthesis of compatible solutes. Compatible solutes refer to low-molecular-weight, highly water-soluble compounds that are typically uncharged, encompassing sugars, polyols, various compounds, amino acids, and their derivatives. In high salinity condition, numerous microorganisms including cyanobacteria re-synthesize and accumulate substantial quantities of small molecular compatible solutes such as sucrose, glucosylglycerol (GG), glycine betaine, etc., to maintain cell turgor pressure and safeguard macromolecular structures without disrupting cellular metabolism^[19,20]^. ii) Active ion transport via specific transporter proteins. Certain ion transport proteins are directly involved in ion transportation, such as the Mrp (Multiple resistance and pH adaptation) system, various Na^+^/H^+^ transport proteins, and Na^+^-ATP synthase^[21–23]^. iii) Regulation of key metabolic pathways. Essential metabolic pathways like the Calvin-Benson-Bassham cycle, glycolysis, and the tricarboxylic acid (TCA) cycle will play pivotal roles in responding to stress^[24]^. Under salt stress, cyanobacteria accumulate lipids and alkanes^[25]^, and their internal chaperone proteins^[26]^, heat shock proteins^[27]^, and antioxidant enzyme activity^[28]^ undergo specific changes. Moreover, they enhance the production of second messengers like the cyclic-di-AMP (c-di-AMP) and cyclic di-GMP (c-di-GMP), which are crucial cellular molecules responding to specific external stimuli and believed to be the initial messengers in environmental adaptation processes^[29,30]^. Despite these molecular mechanisms governing salt tolerance in cyanobacteria obtained from systems biology, comprehensive regulatory networks remain unclear. In addition, although genetic engineering has partially enhanced cyanobacterial salinity resistance to some extent^[31–33]^, it is yet insufficient for obtaining high-salinit tolerant strains.

Confronted with this intricate task of modulating salinity adaptation, adaptive laboratory evolution (ALE) emerges as a potent strategy for bolstering microbial robustness akin to natural selection whilst mitigating ecological hazards^[34]^. In this study, we enhanced salt tolerance of the model cyanobacterium *Synechocystis* sp. PCC 6803 (hereinafter referred to as Syn6803) through ALE methodology, resulting in four evolved strains capable of withstanding 6.5% (m/v) NaCl concentration. Subsequent analysis allowed for phenotype identification within these evolved strains. Whole-genome resequencing of both wild-type and evolved strains revealed eight mutated genes, prompting gene knockout, mutant gene restoration, and overexpression experiments in order to elucidate their roles in conferring salt tolerance upon our strain. Additionally, we harnessed an engineered *slr1753*-expressing strain for seawater treatment, effectively reducing Na^+^ levels therein. Lastly, application of our evolved strain significantly bolstered *Brassica rapa chinensis* growth when cultivated in saline soil compared to untreated controls.

## Results

### Acquisition and Characterization of Halotolerant Syn6803

Prior to conducting ALE of salt on the wild-type strain of Syn6803 (WT), it is imperative to ascertain its baseline salt tolerance. Consequently, we compared its growth under normal as well as 4.0%, 5.0%, and 6.0% (m/v) NaCl conditions. As shown in **Figure 1A**, growth of WT decreased by 37.3% at 96 h under 4.0% NaCl stress, while it significantly declined under 5.0% NaCl stress and maintained almost unchanged under 6.0% NaCl stress. Therefore, we selected 4.0% NaCl stress as the initial condition for WT adaptation. When the OD_750 nm_ of the strain reached approximately 0.2 after 96 h of cultivation, it was deemed to have developed tolerance to the stress condition. Subsequently, the stress level was increased by 0.1% for the next round of adaptation. Finally, after around 856 days’ ALE (125 passages), we obtained and isolated four stable evolution strains (designated ALE-1, ALE-2, ALE-3, and ALE-4) capable of withstanding up to 6.5% NaCl with a remarkable increase in tolerance by up to 62.5% compared to the starting strain (**Figure 1B**). Our findings revealed that under normal conditions, ALE-1, −2, −3 and −4 exhibited comparable growth rates to WT, suggesting no significant growth trade-off of the evolved strains occurred during the ALE. In addition, the four ALE strains demonstrated normal growth with a robust rate in the presence of 6.5% NaCl. While variations in growth were observed among the four evolved strains during logarithmic growth phase, they converged to similar states in stationary phase. In conclusion, we identified four evolution strains with consistent and stable growth phenotypes and selected ALE-3 for further analysis.

**Figure 1.**
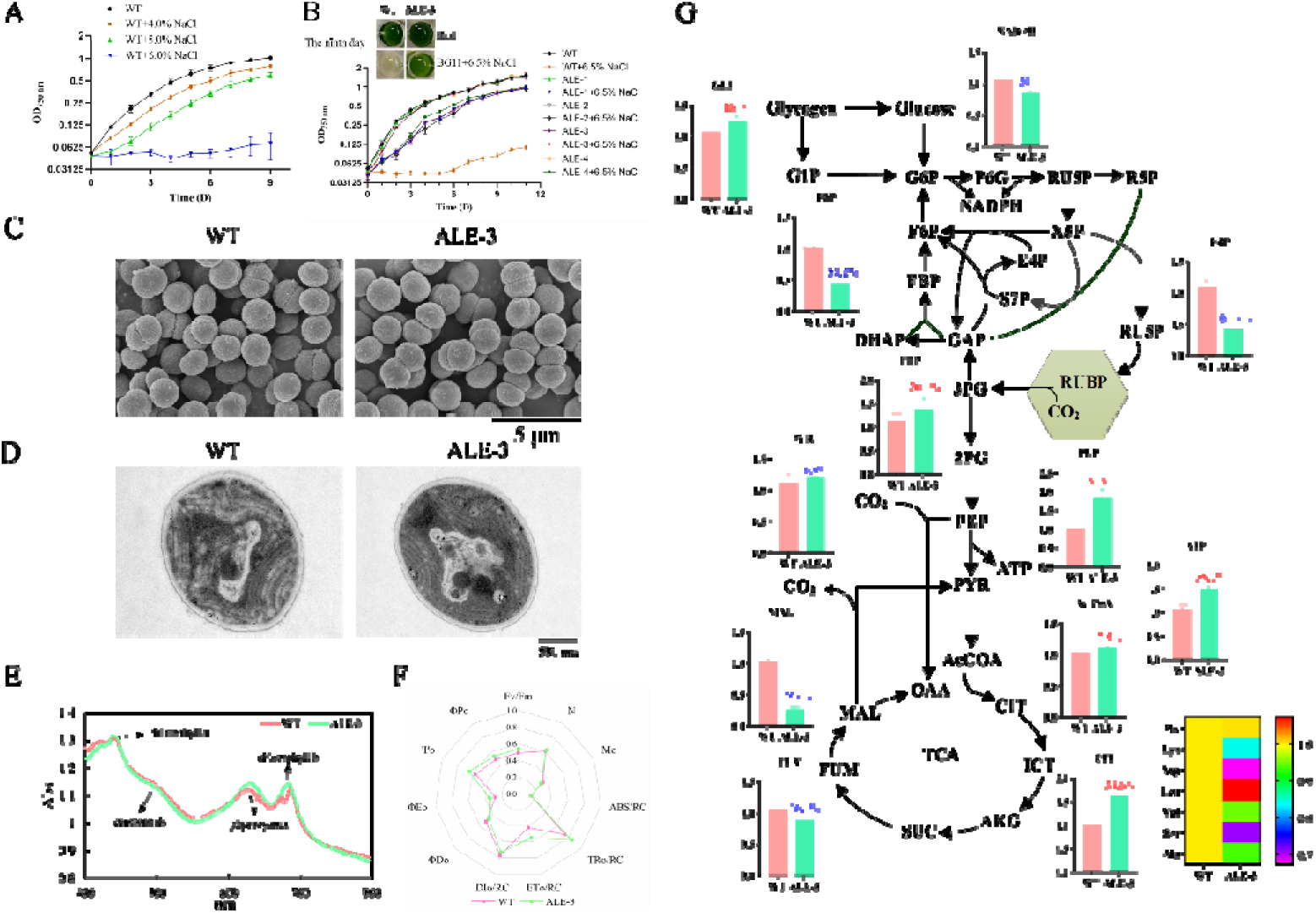
Acquisition and characterization of halotolerant cyanobacteria. (A) Salt tolerance assay of the WT; (B) Comparison of growth between WT and salt-evolved strains under normal and 6.5% NaCl conditions; SEM analysis (C), TEM analysis (D), whole absorption spectrum measurement (E), and chlorophyll fluorescence kinetic analysis (F), Fv/Fm: the maximum photochemical quantum yield of PS□, N: the number of times reaction QA is reduced; Mo: the maximum rate at which reaction QA is reduced, ABS/RC: light energy absorbed by unit reaction center, TRo/RC: energy captured per unit reaction center used to reduce QA, ETo/RC: energy captured per unit reaction center for electron transfer, DIo/RC: energy dissipated per unit reaction center, ΦDo: quantum yield of light energy absorbed by the reaction center used for thermal dissipation, ΦEo: quantum yield of light energy absorbed by the reaction center used for electron transfer, Ψo: the trapped excitation energy drives electron transfer to quantum yield downstream of the QA, ΦPo: the photochemical quantum yield when all reaction centers are open has the same meaning as Fv/Fm, Sm: the reaction QA receptor measures the size of PQ; (G) Metabolomics analysis for both WT and ALE under 4.0% NaCl conditions.

Prolonged exposure to stress may lead to discernible modifications in the cell morphology during the ALE process. Therefore, we employed scanning electron microscopy (SEM) and transmission electron microscopy (TEM) to examine the surface and internal structures of both WT and ALE-3. The findings revealed ALE-3 exhibited no apparent alterations in either morphology or internal structure in comparison to WT (**Figure 1C, 1D**). Further, we evaluated the potential changes in photosynthetic capabilities as extreme salt stress destructed photosystems and led to a bleaching phenotype (**Figure 1B**). Therefore, we measured the full absorption spectrum (400 nm-800 nm) on WT and ALE-3 to assess alterations in their internal light-harvesting pigment contents. Results indicated similar peaks at carotenoids (400 nm-550 nm), chlorophyll a (440 nm), and b (680 nm) for WT and ALE-3 (**Figure 1E**), suggesting comparable carotenoid and chlorophyll contents between them. Nevertheless, ALE-3 exhibited higher phycocyanin content at 625 nm compared to that of WT, suggesting the ALE strain has fully adapted to the high salt stress condition and undergone comprehensive changes in its photosynthesis system. Further, chlorophyll fluorescence kinetic analysis was performed to compare the detail differences of photosynthesis of WT and ALE-3. As shown in **Figure 1F**, the parameters of Fv/Fm, N, Mo, TRo/RC, ETo/RC, ΦDo, ΦEo, Ψo, ΦPo, and Sm in ALE-3 were similar to that of WT, while values of ABS/RC and DIo/RC were higher in ALE-3. This suggested an increased light energy absorption per reaction center and greater energy dissipation as well as a higher quantum efficiency per reaction center in the evolved strain.

To further elucidate the alterations in key intracellular metabolisms, we employed metabolomics analysis for WT and ALE-3 under 4.0% NaCl stress (**Figure 1G**). The following results all represented the change of specific metabolites in ALE-3 compared to that in WT. Notably, the level of glucose (GLU) increased by 16.1% in ALE-3, implying a potential augmentation in energy reserves for its growth. In addition, fructose 1,6-bisphosphate (FBP) and phosphoenolpyruvic acid (PEP) respectively exhibited an increase of 20.1% and 84.9%, possibly indicating heightened carbon flux into the TCA cycle within the evolved strain for biomass accumulation. Conversely, fructose-6-phosphate (F6P) and erythrose-4-phosphate (E4P) experienced reductions by 57.2% and 61.4% correspondingly, suggesting diversion of carbon flux at these sites towards synthesis of other compounds such as compatible solute GG. Under salt stress, the evolved strain can accumulate more ATP than WT due to increased bacterial ATP consumption for ion transport^[35]^. Additionally, amino acid analysis revealed that, except for a slight increase in Leu and Phe, the content of other amino acids was significantly lower in the evolved strain, which may be due to the adjustment of amino acid synthesis to accumulate more ATP in ALE-3.

### Re-sequencing and Identification of Key Genes Conferring Salt Tolerance

To discover the mutations during ALE, we isolated single colonies form ALE-1, −2, −3 and −4 then performed whole-genome re-sequencing. The results revealed base mutations in three genes (*sll1867*, *slr1670*, *sll0689*) and base insertions or deletions in five genes (*slr1753*, *sll1755*, *sll2011*, *sll0377*, *sll0762*) (**Figure 2A**). Next, we investigated the impact of the mutated gene on the strain’s salt tolerance through gene knockout, complementation, and overexpression experiments in WT (**Table 1**). Notably, a neutral gene *slr0168* was also selected for knockout, complementation, or overexpression to make control strain.

**Figure 2.**
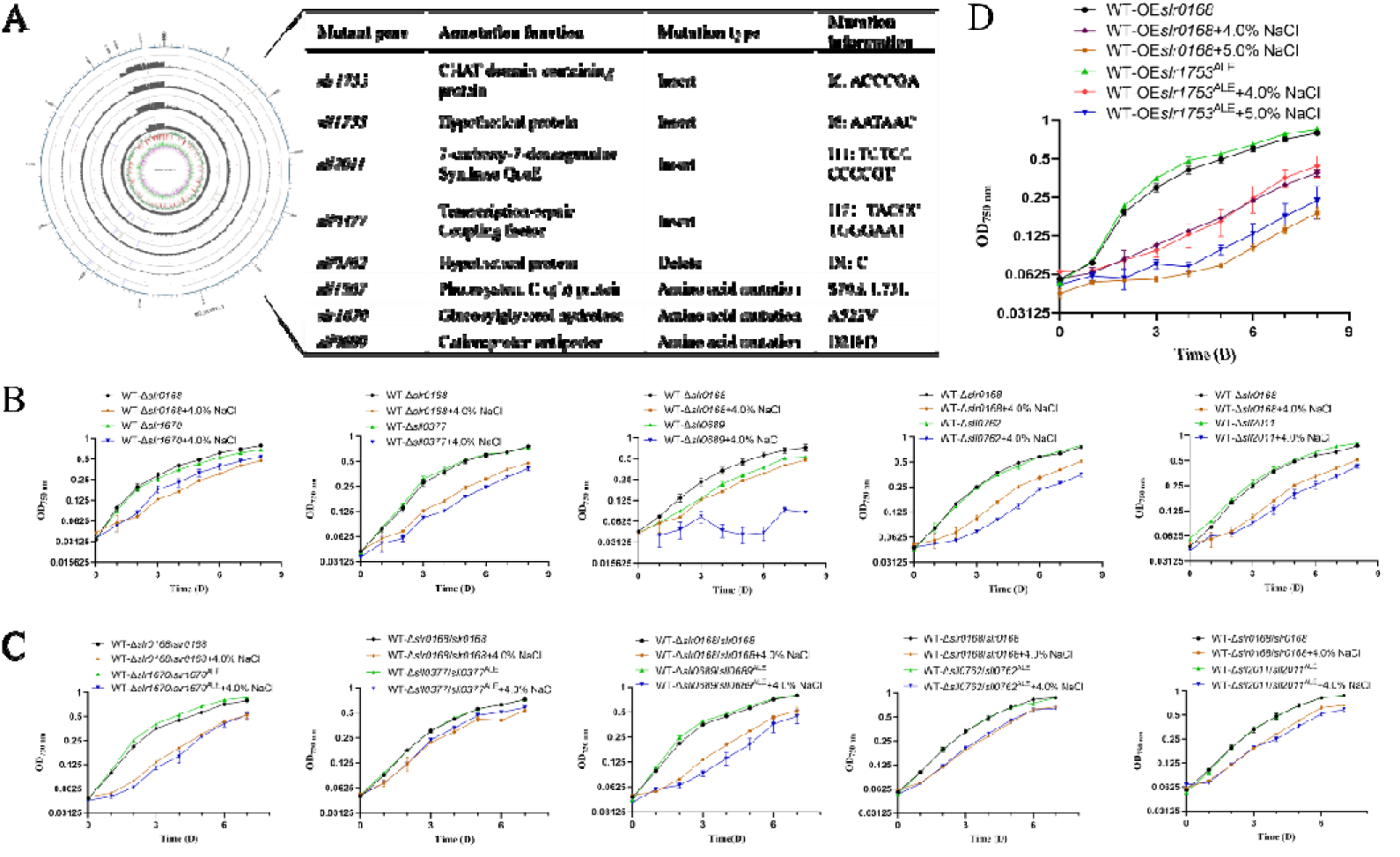
Resequencing of the Genome and Identification of Pivotal Genes Conferring Salt Tolerance. (A) Genome resequencing results and gene mutations of WT and salt-evolved strains; (B) Growth of WT-Δ*slr0168*, WT-Δ*slr1670*, WT-Δ*sll0377*, WT-Δ*sll0689*, WT-Δ*sll0762*, WT-Δ*sll2011* under normal and 4.0% NaCl conditions; (C) Growth of WT-Δ*slr0168*/*slr0168*, WT-Δ*slr1670*/*slr1670*^ALE^, WT-Δ*sll0377*/*sll0377*^ALE^, WT-Δ*sll0689*/*sll0689*^ALE^, ΔWT-*sll0762*/*sll0762*^ALE^, ΔWT-*sll2011*/*sll2011*^ALE^ in normal and 4.0% NaCl conditions; (D) Growth of WT-OE*slr0168* and WT-OE*slr1753*^ALE^ in normal, 4.0%, and 5.0% NaCl conditions; The growth curves were measured in multiple batches, and the growth of the control strains WT-Δ*slr0168*, WT-Δ*slr0168*/*slr0168*, and WT-OE*slr0168* exhibited slight variability across batches.

Via comparing the growth patterns of the gene knockout mutant strains (**Figure 2B**, **Figure S1A**), WT-Δ*sll0689* was found comparatively slower than that of the control strain WT-Δ*slr0168* under normal condition. This phenomenon may suggest the essential role played by *sll0689* in Syn6803, which encodes a Na^+^/H^+^ antiporter crucial for maintaining internal ion homeostasis^[36]^. Under 4.0% NaCl conditions, WT-Δ*slr1753*, WT-Δ*sll1755* and WT-Δ*sll1867* exhibited similar growth trends as WT-Δ*slr0168*. However, distinct variations were evident for WT-Δ*sll2011*, WT-Δ*sll0377*, WT-Δ*sll0762*, WT-Δ*slr1670*, and particularly for WT-Δ*sll0689* compared to WT-Δ*slr0168*, suggesting potential related roles of these genes with salt stress. Among them, WT-Δ*slr1670* exhibited a superior growth phenotype, suggesting its deletion conferred a beneficial effect on salt tolerance. The *slr1670* gene is postulated to possess GG lyase activity, and thus its deletion would lead to the accumulation of more compatible solute GG in the strain, thereby enhancing its salt tolerance^[37]^. Conversely, the respective deletion of *sll2011*, *sll0377*, *sll0762*, and *sll0689* resulted in an inferior growth phenotype compared to WT-Δ*slr0168*, indicating their absence was detrimental to salt tolerance. Among these genes, *sll0377* encodes a transcription-repair coupling factor—a widely conserved bacterial protein involved in mediating transcription-coupled DNA repair^[38]^—suggesting that its deletion may impact DNA repair processes and consequently decrease salt tolerance. In addition, *sll0689* encodes a Na^+^/H^+^ antiporter protein essential for enabling growth under salt stress when present but causing inability to grow under salt stress upon deletion^[36]^. Moreover, *sll2011* encodes the 7-carboxyl-7-deazaguanine synthetase QueE with unclear implications for bacterial salt tolerance, while the function of *sll0762* remains unknown. Further studies are required to elucidate the roles of these genes in salt tolerance. Through gene complementation using mutated sequences form the ALE strains, the growth patterns of the 8 complementation strains under salt stress closely resembled those of the control strain WT-Δ*slr0168*/*slr0168* (**Figure 2C**, **Figure S1B**), suggesting the salt-responsive phenotype can be mitigated. However, no strain exhibited enhanced salt tolerance. Further, direct overexpression of the mutated gene was performed at a neutral locus *slr0168* in WT. WT-OE*sll0689*^ALE^ was not obtained whose overexpression may disrupt intracellular ion balance. Though no significant differences in growth were observed under 4.0% NaCl stress, WT-OE*slr1753*^ALE^ exhibited noticeable growth advantage under 5.0% NaCl (**Figure 2D**, **Figure S1C**). It has been reported that the Slr1753 protein is located on the cell membrane^[39]^. Therefore, we hypothesize that overexpression of *slr1753* may alter the composition of the cell membrane and influenced ion transport, thereby enhancing salt tolerance.

### Functional Analysis of Genes Associated with Salt Tolerance

Growth patterns have demonstrated that deletion of *slr1670* and overexpression of mutated *slr1753* could improve the salt tolerance of Syn6803. Notably, the phenotype of WT-Δ*slr1670* became more significantly when we assessed the growth condition in 5.0% NaCl (**Figure 3A**). Given that *slr1670* encodes a GG hydrolase (**Figure 3B**), we further quantified the GG content in WT-Δ*slr1670* and WT-Δ*slr0168* under 5.0% NaCl condition. As expected, a 266.3% increase of intracellular GG content was observed in WT-Δ*slr1670* compared to that in WT-Δ*slr0168*, with no detectable extracellular GG (**Figure 3C**). Consistently, we found the intracellular GG content of ALE-3 was 82.7% higher than that of WT under 5.0% NaCl condition (**Figure 3D**, **E**), suggesting the elevated intracellular GG content caused by mutated *slr1670* contributed to the enhanced salt tolerance.

**Figure 3.**
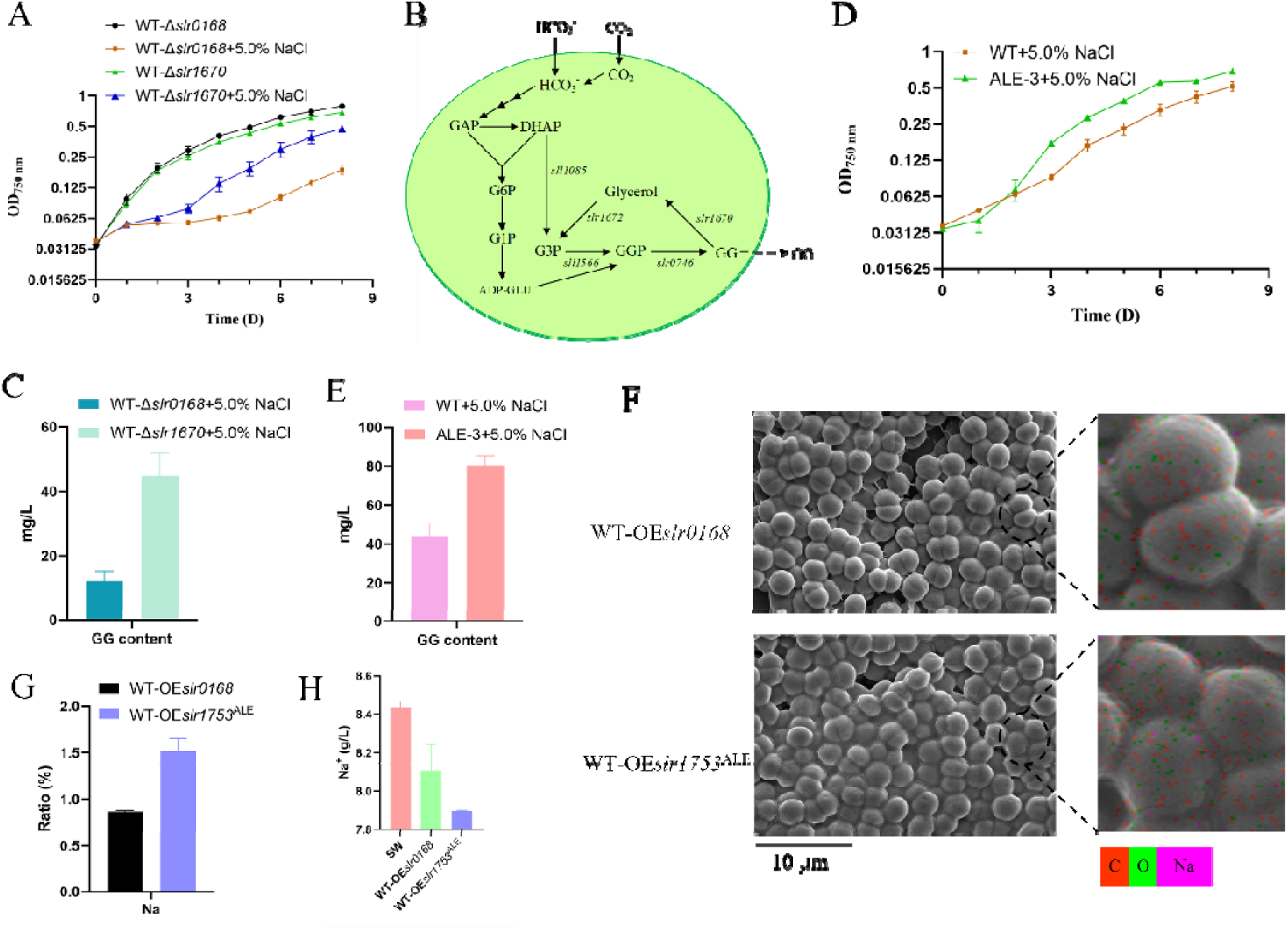
Functional analysis of genes associated with salt tolerance. (A) Growth of WT-Δ*slr0168* and WT-Δ*slr1670* under normal and 5.0% NaCl conditions; (B) Synthesis pathway of GG in Syn6803; (C) Intracellular GG production of WT-Δ*slr0168* and WT-Δ*slr1670* under 5.0% NaCl conditions; (D) Growth of WT and ALE under 5.0% NaCl conditions; (E) Intracellular GG production of WT and ALE under 5.0% NaCl conditions; (F) SEM EDX analysis of cell surfaces for WT-OE*slr0168* and WT-OE*slr1753*^ALE^ strains; (G) Sodium element content on the cell surface of WT-OE*slr0168* and WT-OE*slr1753*^ALE^ strains, as determined by EDX analysis; (H) The intracellular ROS content of WT-OE*slr0168* and WT-OE*slr1753*^ALE^ under normal and 5.0% NaCl culture conditions; (I) The residual Na^+^ content following treatment of WT-OE*slr0168* and WT-OE*slr1753*^ALE^ with seawater.

As Slr1753 is possibly located on the cell membrane^[39]^, we speculated it may be a transporter protein related with ion transport. Thus, SEM energy dispersive X-ray (EDX) analysis was utilized to compare the surface ion differences between WT-OE*slr0168* and WT-OE*slr1753*^ALE^. As a result, the WT-OE*slr1753*^ALE^ strain exhibited increased adsorption of Na on its surface (**Figure 3F, 3G**, and **Figure S2**), which demonstrated our hypothesis. Encouraged by the results, the Na^+^ ion adsorption capability of WT-OE*slr1753*^ALE^ in simulated seawater desalination was investigated. After collecting 10 OD_750 nm_ strains and subjecting them to seawater treatment for 72 hours, the sodium ion content was quantified using ICP. The results revealed a reduction of 6.35% in the Na^+^ ion content by WT-OE*slr1753*^ALE^ (**Figure 3I**). In conclusion, leveraging the effective Na^+^ ion adsorption capacity of this strain presents a novel approach towards seawater desalination.

### Utilization of ALE strains in Bioremediation of Saline-Alkaline Soil

The management of soil salinization typically includes implementing drainage systems to remove excess salts, enhancing soil structure through improved drainage, and occasionally incorporating amendments to enhance fertility. The introduction of salt-tolerant cyanobacteria has been considered a promising approach for ameliorating saline soils^[40]^. Thus, we aimed to evaluate if our evolved strain would exhibit superior properties in bioremediation of saline-alkaline soil. Detailly, we introduced WT and ALE-3 into low- (total salt of 2.5 mg/kg soil) and high concentration (total salt of 11.0 mg/kg soil) saline-alkali soil during cultivation of *Brassica rapa chinensis*. The plant culture groups were designated as O-BC for those without the strain, WT-BC for those with the WT, and ALE-BC for those with ALE-3. In low-concentration saline-alkali soil, *Brassica rapa chinensis* exhibited accelerated growth compared to that in high-concentration one (**Figure 4A, 4B**). We analyzed and determined its germination rate, height, as well as soil TOC, total salt, and pH after one week of cultivation in a controlled environment (**Figure 4C, 4E**, and **Figure S3**). Addition of WT and ALE-3 to low-concentration saline-alkali soil was found to increase the germination rate of *Brassica rapa chinensis* by 5.3% and 10.6%, respectively, with no significant difference in average height. While WT did not significantly affect soil TOC, it led to a reduction in total salt and pH by 35.2% and 0.036%, respectively; ALE-3 increased soil TOC by 1.53% and reduced total salt and pH by 32.9% and 0.47%, respectively. Conversely, in high-concentration saline-alkali soil, the growth rate was slower. We analyzed and determined its various indicators after three weeks of cultivation in the controlled environment with additional bacterial solution administered every seven days (**Figure 4B**). It was observed that the addition of WT increased the germination rate and average height of *Brassica rapa chinensis* by 89.4% and 35.8%, respectively, in high salt-alkali soil, and the application of ALE-3 led to a remarkable increase of 184.2% and 43.8%, respectively. Furthermore, the addition of WT resulted in an 11.7% increase in soil TOC and a decrease in soil pH by 1.76%, while it did not reduce the total salt content of the soil. Conversely, the application of ALE-3 led to a substantial increase in soil TOC by 25.3% and reductions in both total salt content by 1.82% and pH by 1.91% (**Figure 4D** and **4F).** The aforementioned evidence indicates that cyanobacteria can enhance soil fertility and promote plant growth, suggesting that cyanobacteria with higher salt tolerance have greater potential for improving saline soils.

**Figure 4.**
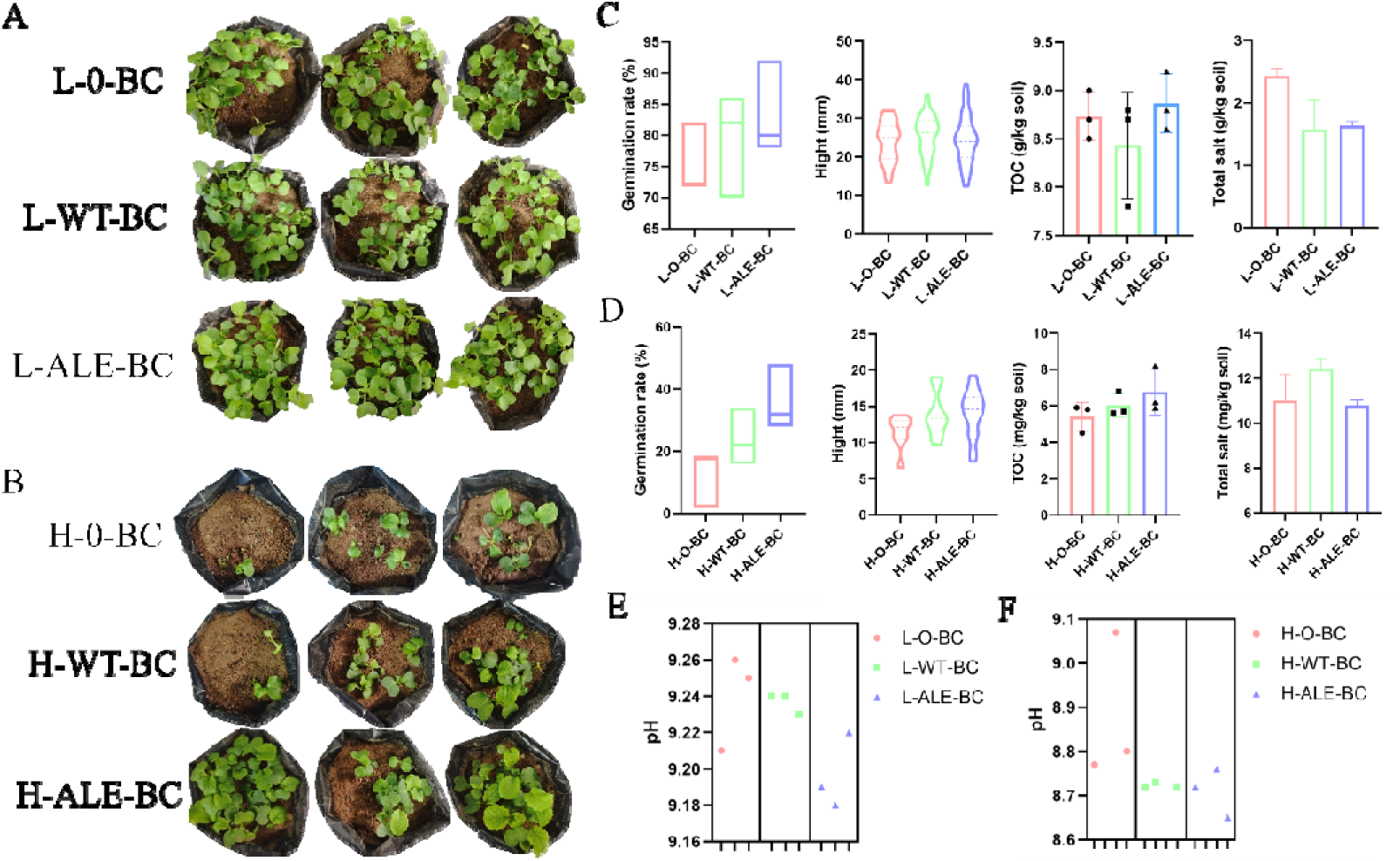
Utilization of evolved cyanobacteria in saline-alkaline soil. (A) L-O-BC, L-WT-BC, and L-ALE-BC denote the growth conditions of *Brassica rapa chinensis* in low-concentration saline-alkali soil without inoculation, with inoculation of the wild-type Syn6803, and with inoculation of salt-evolved Syn6803, respectively; (B) H-O-BC, H-WT-BC, and H-ALE-BC denote the growth conditions of *Brassica rapa chinensis* in high-concentration saline-alkali soil without inoculation, with inoculation of the wild-type Syn6803, and with inoculation of salt-evolved Syn6803, respectively; After cultivating *Brassica rapa chinensis* in low-concentration saline-alkali soil for a week, the germination rate, plant height, soil TOC, total salt content (C), and pH (E) were subjected to statistical analysis; Following three weeks of cultivation in high-concentration salt-alkali soil, the germination rate, plant height, soil TOC, total salt content (D), and pH (F) were statistically analyzed.

## Discussion

With the recent utilization of transgenic cyanobacteria as a platform for producing various biofuels and chemicals^[41–43]^, coupled with the limited freshwater supply on Earth, large-scale cultivation of cyanobacteria will inevitably shift to seawater in the future. Therefore, it is imperative to comprehensively elucidate the salt-tolerance mechanisms of cyanobacteria in order to develop salt-tolerant freshwater cyanobacterial platforms. This study utilized ALE technique to enhance the salt tolerance of Syn6-803, enabling the strain to withstand 6.5% NaCl, a feat not previously achieved through genetic modification or other means^[20,44,45]^.

Upon further analysis, it was determined that *slr1670*, involved in GG synthesis, and *slr1753*, associated with Na^+^ adsorption, are pivotal genes contributing to the enhanced salt tolerance of the evolved strain. Quantitative measurements revealed that the salt-evolved strain exhibits increased GG synthesis. Given its low sweetness, high water retention capacity, protective properties for large molecules, and anti-tumor activity, GG demonstrates significant potential for applications in cosmetics, food service, and pharmaceuticals^[46]^. Consequently, there is potential to develop it as a platform for GG production in the future. Additionally, to cope with the escalating water demand driven by factors such as population growth, industrial expansion, tourism development, and agricultural progress, numerous regions are turning to seawater desalination as a means of augmenting water supply. Technological advancements and heightened environmental consciousness have led to the widespread adoption of seawater desalination techniques including reverse osmosis and distillation processes for converting saline water into freshwater^[47,48]^. Additionally, solar-driven evaporation and condensation systems, ion exchange membrane technology, and other innovations are being employed to secure additional freshwater resources^[49]^. Therefore, considering WT-OE*slr1753*^ALE^’s Na^+^ adsorption capacity offers a fresh perspective and potential solution for addressing the global freshwater scarcity issue.

Soil salinization represents a critical ecological safety concern, and the utilization of cyanobacteria for saline soil remediation constitutes an innovative approach. Firstly, cyanobacteria can absorb and fix excessive salts in the soil, thereby reducing salt content; additionally, the exopolysaccharides they produce can improve soil aggregate structure and water retention capacity^[17]^. Secondly, the decomposition of cyanobacterial biomass contributes organic matter to the soil, thus improving its fertility^[16]^. Finally, cyanobacteria can stimulate the growth of beneficial microbial communities in the soil while also promoting plant growth through their production of plant hormones^[50]^—ultimately achieving biological remediation. Similarly, during the cultivation of *Brassica rapa chinensis*, we observed that the evolved strain exhibited remarkable efficacy in the biological reclamation of saline-alkali soil.

In the future, we aim to further augment the salt tolerance of the strain through accelerated evolutionary methods, investigate its enduring effects and influence on saline soil, and elucidate the mechanism of Na^+^ adsorption by Slr1753 via protein structure analysis, among other strategies.

## Conclusion

In this study, we enhanced the salt tolerance of cyanobacteria through ALE. Phenotypic analysis revealed that the salt-evolved bacteria exhibited full adaptation to high-salt environments, maintaining similar morphology and photosynthesis characteristics to the wild type while undergoing significant changes in central metabolism under salt stress. Experimental analysis of mutated genes indicated that at least five genes were associated with the enhanced salt resistance of the evolved bacteria, but only gene *slr1753* showed significant effects when overexpressed individually, suggesting potential synergistic interactions among multiple genes. Subsequent analysis identified *slr1670* and *slr1753* as the two key genes responsible for enhanced salt resistance. Moreover, the adapted bacteria show promise for treating salt-alkali areas, with initial results indicating positive effects. Continuous exploration and optimization of this technology may lead to more effective, sustainable, and environmentally friendly approaches for addressing salt-alkali areas, thereby contributing to agricultural production and ecological protection

## Materials and methods

### Cultivation and Construction of Cyanobacterial Strains

The wild-type (WT), salt-evolved (ALE), and engineered strains of Syn6803 in this investigation were cultivated in BG11 medium (pH 7.5) under a light intensity of 50 μmol photons m^−2^ s^−1^ in an illuminating incubator at 30 °C with agitation at 160 rpm (HNY-211B Illuminating Shaker, Honour, Tianjin, China). Cell density was assessed using an ELx808 Absorbance Microplate Reader (BioTek, VT, USA) at OD_750 nm_. Cultivate the wild-type, salt-evolved, and engineered strains of Syn6803 in a shaking incubator for pre-culture. When the pre-cultured strains reach the logarithmic phase, take 1 OD of the cells and resuspend them in 25 mL of normal saline BG11 liquid culture medium to achieve an initial OD_750 nm_ of 0.04. Each cultivation condition includes three biological replicates. Every 24 hours, transfer 200 μL of the cell suspension to a 96-well plate for measurement of OD_750 nm_ using an enzyme-linked immunosorbent assay (ELISA) reader.

The gene knockout strains were generated by amplifying the chloramphenicol resistance cassette and fusing it with DNA fragments upstream and downstream of the target gene, resulting in a linear DNA fragment. This linear fusion fragment was directly utilized for gene knockout in Syn6803 through natural transformation to obtain chloramphenicol-resistant transformants. The chloramphenicol-resistant transformants were subsequently cultured multiple times in BG11 medium supplemented with 50 μg/mL chloramphenicol to achieve complete segregation. To complement the knocked-out gene in the mutant, a mutated gene from the salt-evolved strain, along with the streptomycin resistance cassette and DNA fragments upstream and downstream of the knocked-out gene, were amplified and fused into a linear DNA fragment. This linear fusion fragment was then introduced into the knockout mutant strain via natural transformation. Spectinomycin-resistant transformants were selected and cultured several times in BG11 medium supplemented with 50 μg/mL spectinomycin to achieve complete separation. The overexpression strain construct was created by fusing the mutated gene from the salt-evolved strain with a plasmid scaffold containing upstream and downstream gene sequences of *slr0168* as well as *cpc560* promoter. The constructed plasmid was introduced into Syn6803 through natural transformation, followed by selection of chloramphenicol-resistant transformants. These strains underwent multiple rounds of culturing in BG11 medium supplemented with 50 μg/mL chloramphenicol to ensure complete segregation. Mutant strains were confirmed using colony PCR and sequencing analysis.

### Electron Microscope Analysis

Zhongke Baitest Technology Co., Ltd. (Beijing, China) offers transmission electron microscope (TEM) and scanning electron microscope (SEM) analysis services. For sample preparation, cells in logarithmic growth phase are collected by centrifugation and washed twice with PBS buffer (pH 7.0; 0.1 M). Subsequently, a 2.5% glutaraldehyde solution is slowly added to the cell precipitate at 4 °C for 12 hours. For TEM analysis, the glutaraldehyde solution is removed and the sample is washed three times with PBS buffer before undergoing fixation steps including osmium tetroxide treatment, dehydration using an ethanol gradient, infiltration with acetone-resin mixture, and gradual heating in an oven. Thin sections of the sample (70-90 nanometers) are then obtained using a UC7 ultramicrotome (Leica, Germany), followed by staining with uranyl acetate and lead acetate prior to observation on a JEOL JEM-1200EX transmission electron microscope (JEOL, Tokyo). In SEM analysis, after removal of the osmium tetroxide solution and triple washing with PBS buffer, the sample undergoes dehydration using an ethanol gradient before being dried for observation on a Hitachi SU8020 SEM instrument (Hitachi, Tokyo). Elemental automatic qualitative analysis is conducted through integration of an EX-250 energy dispersive X-ray spectrometer (Horiba, Kyoto) with the SEM^[51]^.

### Measurement of Absorption Spectrum and Photophysiological Parameters

Cultivate Syn6803WT and ALE in 4.0% NaCl until reaching the logarithmic growth phase, standardize the cell density using OD_750 nm_, and measure it with a UV-1601 spectrophotometer (Beijing Beifeng Ruili Instrument Co., Ltd., China). Analyze the data using UVprobe software (version X). For measuring Chlorophyll fluorescence kinetics, first dark-adapt the logarithmic phase strain for 15 minutes, then transfer 3 ml of the culture to an AquaPen AP 110/C handheld PAM fluorimeter (Drasov, Czech Republic) for measurement.

### ROS Measurement

After culturing WT-OE*slr0168* and WT-OE*slr1753*^ALE^ for 4 days, bacterial samples were collected (volume × OD_750 nm_ = 0.1), and the Reactive Oxygen Species Assay Kit (Shanghai Binyuntian Biotechnology Co., Ltd.) was used for ROS determination. Briefly, the fluorescent probe DCFH-DA, which is non-fluorescent itself, can freely permeate the cell membrane and enter the cell. Upon hydrolysis by intracellular esterase, DCFH is produced. Intracellular reactive oxygen species oxidize non-fluorescent DCFH to generate fluorescent DCF. The fluorescence of DCF was measured using an Infinite E Plex Microplate Reader (TECAN, Switzerland) at EX 488 nm and EM 525 nm.

### Extraction and Detection of Intracellular Metabolites

After culturing WT and ALE strains in 4.0% NaCl for 4 days, bacterial samples (volume × OD_750 nm_ = 8) were collected. The bacterial cells were suspended in a solution of 80% methanol/H_2_O (8:2, v/v), followed by repeated freeze-thawing in liquid nitrogen for five cycles. The supernatant was obtained through centrifugation. Subsequently, the bacterial cells from the previous step were re-suspended in an 80% methanol/H_2_O solution and subjected to repeated freeze-thawing in liquid nitrogen. The combined supernatants from both steps were lyophilized (ZLS-1, Her-exi, Hunan, China). and then reconstituted with deionized water before filtration and analysis. The detection method employed in this study closely resembled that of the reference laboratory^[31,52]^, utilizing liquid chromatography-quadrupole quadrupole mass spectrometry (LC-QQQ-MS) for metabolite detection. The XBridge Amide column (Waters, Milford, MA, USA, 150 × 2.1 mm, 3.5 μm) was selected and operated in MRM mode. The data underwent processing using Qualitative Analysis B.0.

### Extraction and detection of intracellular and extracellular GG content

The determination method was adapted from previous studies and subjected to slight modifications^[53,54]^. Following a 5-day culture period, 1 ml of the culture liquid was extracted and centrifuged at 12,000×g for 5 minutes to isolate the supernatant. Subsequently, the supernatant was filtered through a 0.22 μm filter to determine the extracellular GG content. The cell pellet was reconstituted in an 80% ethanol solution and incubated at 65 LJ for 4 hours. Afterward, the cell pellet underwent centrifugation at 12,000 rpm for 5 minutes; the resulting supernatant was then evaporated using a rotary evaporator and dissolved in 1 ml deionized water. The cellular GG content was further assessed by filtration through a 0.22 μm filter. Subsequently, the GG sample underwent analysis using an Agilent 1260 Series Binary High-Performance Liquid Chromatography System (Agilent Technologies, California, USA) equipped with an Am inex HP X-87C column (300×7.8 mm, 9μm; Bio-Rad, California, USA). The column was maintained at 80°C and the water flow rate was set to 0.6 mL/min. The identical method was employed for analyzing various concentrations of GG standard (GG standard provided by Qingdao Zhongkezhiqing Biotechnology Development Co., Ltd.), resulting in the generation of a standard curve.

### Quantification of Na^+^

Inoculate the cyanobacteria strain (volume × OD_750 nm_ = 10) into saltwater or seawater (Shandong Yantai, China, 121°17’17.052’ E, 37°34’56.075’ N) and incubate it on a shaker for 3 days. Extract 5 ml of the culture supernatant, subject it to bacterial filtration, and dispatch the sample to Zhongke Baitest Technology Co., Ltd. (Beijing, China) for analysis. The Na^+^ content in the sample was quantitatively determined using an iCAP 7400 ICP-OES analyzer (ICP-OES; Thermo Fisher, Carlsbad, CA, USA).

#### Brassica rapa chinensis cultivation

After low- or high-concentration saline-alkali soil has been thoroughly dried at high temperature and sieved through a 10-mesh 2mm sieve, it is mixed with vermiculite (2:1, v/v). Subsequently, 200g of the mixture is placed in a flowerpot, followed by the addition of 100ml of water and thorough stirring. Upon adding 50 seeds of *Brassica rapa chinensis* to the pot, the soil is covered. A cyanobacteria solution (volume × OD_750 nm_ = 1) is then sprayed on the soil surface before placing it in a plant growth chamber under conditions of 25°C and a light-dark cycle of 16/8h. Regular watering is carried out throughout the cultivation period. The germination rate and height of *Brassica rapa chinensis* are statistically recorded one week after cultivation for low concentration saline-alkali soil conditions, while for high concentration saline-alkali soil conditions, these parameters are recorded three weeks after cultivation. Additionally, cyanobacteria solution is added every seven days during the cultivation period.

### Soil Analysis

After conducting statistical analysis on the germination rate and height of *Brassica rapa chinensis*, the soil was extracted from the pots and sent to Zhongke Baitest Technology Co., Ltd. (Beijing, China) for soil analysis. The total organic carbon (TOC) was determined using the potassium dichromate oxidation-external heating method and a digital titrator (DE-M); pH was measured using the electrode method and a pH meter (PHS-3Cb); and total salt content was assessed using the mass method with a one-thousandth electronic balance (FA1004).

## Acknowledgments

This research was supported by grants from the Ministry of Ecology and Environment Key Laboratory on Biodiversity and Biosafety, the National Key Research and Development Program of China (Grant no. 2020YFA0906800), the National Natural Science Foundation of China (Grant nos. 32371486 and 32270091), the Natural Science Foundation of Tianjin (Grant no. 23JCYBJC01680), and the Fundamental Research Funds of CAF (project No. CAFYBB2021MB005).

