## Supplementary figures and images for "Enhanced Salt Tolerance in *Synechocystis* sp. PCC 6803 Through Adaptive Evolution: Mechanisms and Applications for Environmental Bioremediation"

### Supplemental figures

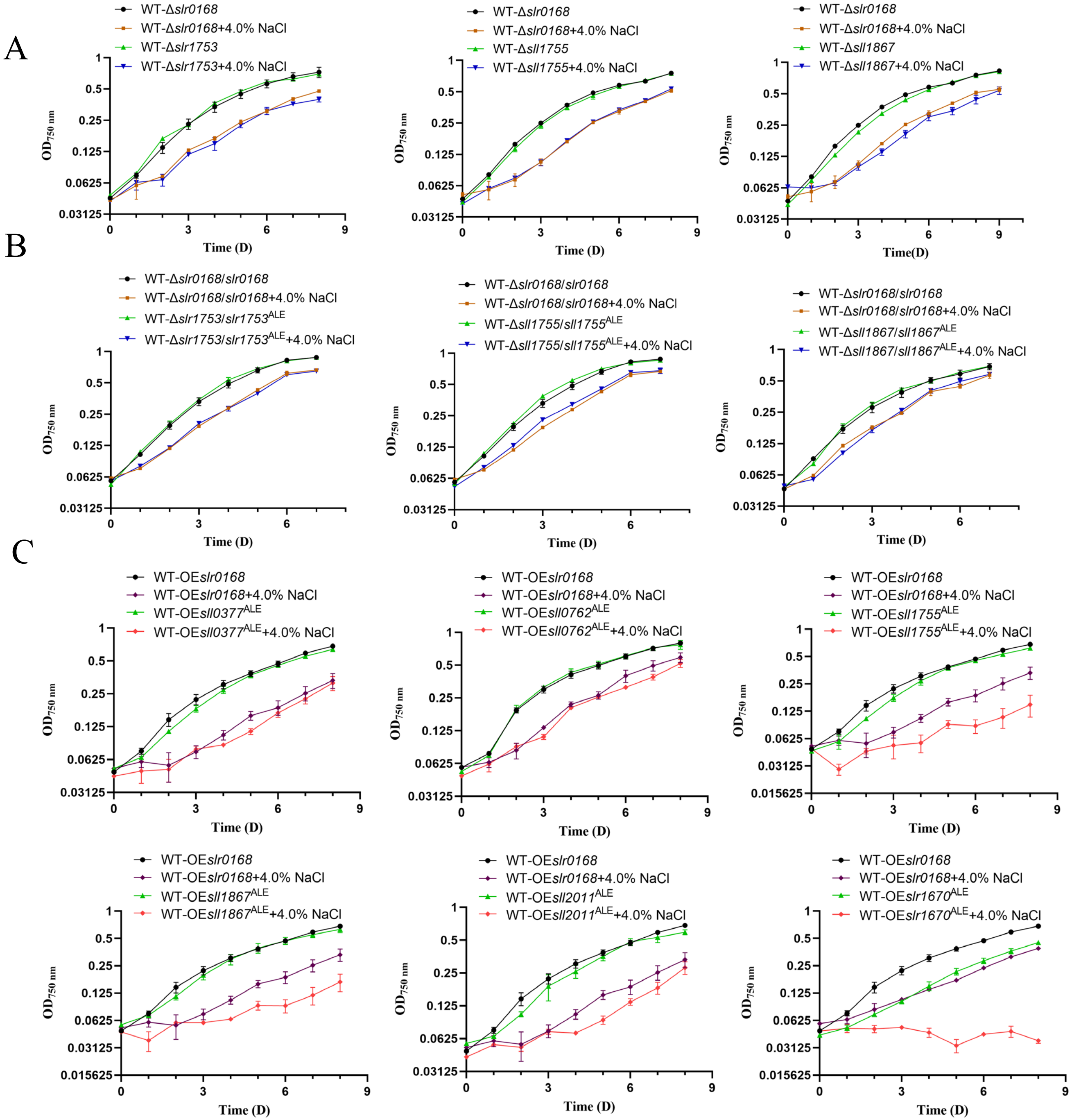

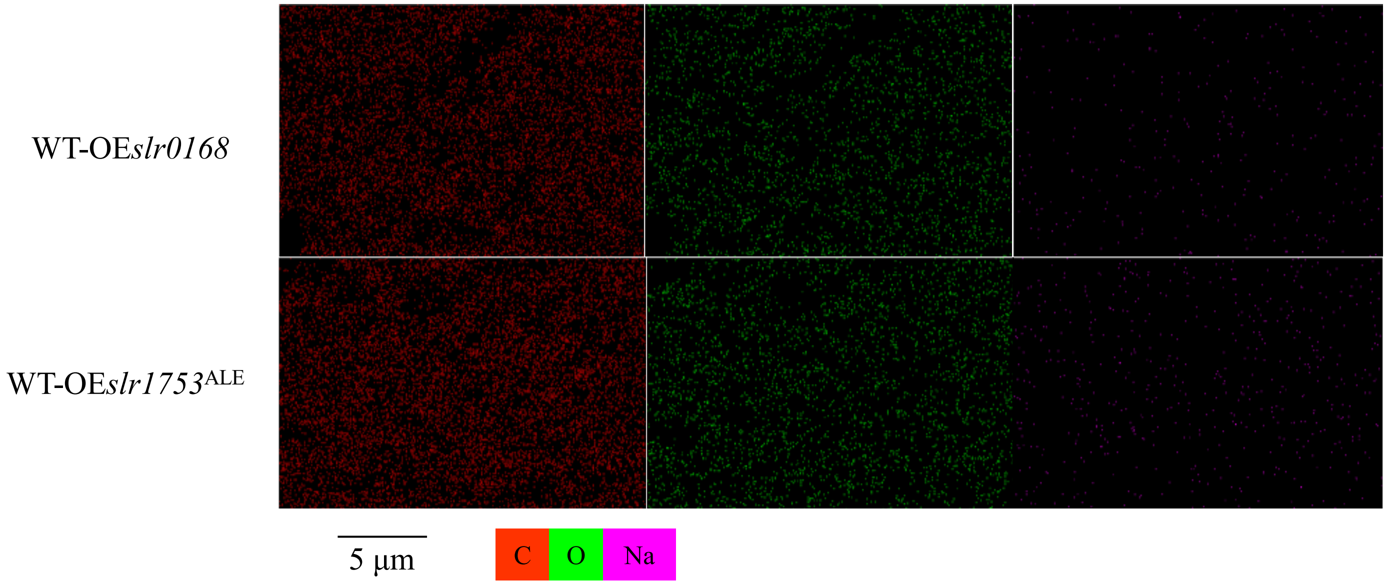


Figure S2

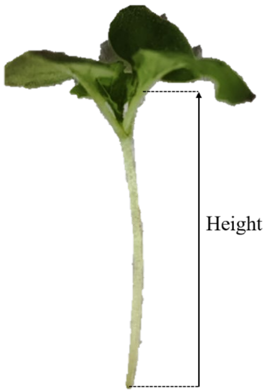


Figure S3
